# Optimizing automated detection of high frequency oscillations using visual markings does not improve SOZ localization

**DOI:** 10.1101/2020.09.14.297309

**Authors:** Trisha Mendoza, Casey L. Trevino, Daniel W. Shrey, Jack J. Lin, Indranil Sen-Gupta, Beth A. Lopour

## Abstract

**Objective:** High frequency oscillations (HFOs) are a biomarker of the seizure onset zone (SOZ) and can be visually or automatically detected. In theory, one can optimize an automated algorithm’s parameters to maximize SOZ localization accuracy; however, there is no consensus on whether or how this should be done. Therefore, we optimized an automated detector using visually identified HFOs and evaluated the impact on SOZ localization accuracy.

**Methods:** We detected HFOs in intracranial EEG from 20 patients with refractory epilepsy from two centers using (1) unoptimized automated detection, (2) visual identification, and (3) automated detection optimized to match visually detected HFOs.

**Results:** SOZ localization accuracy based on HFO rate was not significantly different between the three methods. Across patients, visually optimized detector settings varied, and no single set of settings produced universally accurate SOZ localization. Exploratory analysis suggests that, for many patients, detection settings exist that would improve SOZ localization.

**Conclusions:** SOZ localization accuracy was similar for all three methods, was not improved by visually optimizing detector settings, and may benefit from patient-specific parameter optimization.

**Significance:** Visual HFO marking is laborious, and optimizing automated detection using visual markings does not improve localization accuracy. New patient-specific detector optimization methods are needed.

## 1. Introduction

For over two decades, the link between high frequency oscillations (HFOs) and seizure generating brain regions has been extensively studied [1, 2]. Interictal HFOs have a demonstrated association with the seizure onset zone (SOZ), as they occur more frequently in the intracranial EEG (iEEG) channels in which seizure activity is first visible [3, 4]. Moreover, the removal of HFO-generating regions has been correlated with favorable post-surgical outcomes [5–7]. However, many of these studies rely on group level analyses; for individual patients, identification of HFO-generating regions is not yet reliable enough to be used as a tool for surgical planning, as post-surgical outcomes cannot be accurately predicted [8, 9].

One strategy for increasing the accuracy of HFOs as a predictive biomarker of surgical outcome has been to improve the methods used for detection [10]. Because manually marking HFOs is a laborious task with low interrater reliability [11, 12], researchers have developed numerous fast and objective algorithms for automatic HFO detection [10, 13]. While use of these automatic detectors is becoming more widespread, their implementation varies significantly across studies, as no consensus has been reached on how to properly select parameter values for a given algorithm or for a specific dataset. These settings may include parameters such as an amplitude threshold, a minimum duration, or a minimum number of oscillations for each event. The combination of the selected parameters directly impacts the characteristics and the rate (number per minute) of the detected HFOs.

One common approach for implementation of automated detection is to use the settings from the original publication, which allows researchers to compare results between studies and further validate the performance of the detector. It is typical to apply the same detector settings to all patients in a study [8, 14-16]. However, the application of an existing HFO detection algorithm to a new dataset often results in lower detection accuracy compared to the original application [15, 17-19]. This decrease in accuracy can be attributed to variations in data characteristics, differences in electrode types, or analysis of disparate brain regions [20]. Moreover, it is common to find that standardized detector settings exhibit favorable performance on the group level, while localization accuracy for a subset of patients is poor [8]. One strategy to overcome this limitation and improve localization accuracy is to customize detection parameters. In many cases, parameters are chosen such that detected events closely match visually identified HFOs, either from a subset of the patient population [8, 15, 16] or a subset of channels in each patient [21]. Other times, specific settings are selected because they have reported to outperform the original ones [17]. However, it is unknown if implementing these forms of detector optimization results in improved SOZ localization. A systematic examination of optimization strategies is needed to assess the value of patient-specific customization, as is characterization of the relationship between SOZ localization accuracy and HFO detector settings.

Therefore, the objectives of this study were to test the impact of HFO detector optimization on SOZ localization accuracy and to assess the variability of the accuracy across different detector settings in individual subjects. To do this, we performed automatic and visual HFO detection on iEEG data from two centers. We calculated SOZ localization accuracy based on the HFO rates generated in three ways: (1) Using the unoptimized automatic HFO detection settings from the original publication, (2) Using visual detection only, and (3) Using automatic HFO detection with settings optimized to match visual detection. The relative performance for each approach sheds light on the utility of detector optimization and the need for patient-specific approaches.

## 2. Methods

### 2.1 Clinical data collection

Twenty patients with refractory epilepsy from two medical centers were identified retrospectively and included in this study. Seven of these patients underwent implantation of intracranial electrodes for presurgical evaluation from June 2015 to March 2017 at the University of California, Irvine (UCI) Medical Center. We will refer to this as the UCI dataset. We included only patients with a postoperative outcome of Engel Class I, which suggests that the clinical SOZ localization was successful. For each patient, the SOZ was defined as electrodes involved within the first three seconds of seizure onset, based on visual assessment of the ictal iEEG. If patients had multiple seizures with different regions of onset during recording, all SOZ channels were considered SOZ. The demographic and clinical characteristics of these patients are shown in Table 1. Collection and analysis of retrospective patient data for this study was approved by the Institutional Review Board of the University of California, Irvine.

**Table 1.** Patient demographics of the UCI dataset. Abbreviations: AM = amygdala; AH = anterior hippocampus; depth = depth electrode; FLE = frontal lobe epilepsy; grid = subdural grid electrode; HH = head of hippocampus; HP = hippocampus; L = left; MF = mesial frontal; PFC = pre-frontal cortex; TH = tail of hippocampus; TLE = temporal lobe epilepsy; R = right

| Patient | Age, Gender | Seizure semiology | MRI | Diagnosis | Electrodes | SOZ channels | Surgery | Engel outcome | Postoperative follow-up (months) |
| --- | --- | --- | --- | --- | --- | --- | --- | --- | --- |
| 1 | 44, M | Focal Impaired Awareness Seizures | Cortical Dysplasia in PFC | FLE | 1 grid 8x8<br>2 grid 2x8<br>1 grid 4x8 | RMF28-32 | R Frontal Lobectomy | IA | 47 |
| 2 | 50, M | Focal Impaired Awareness Seizures | White matter changes in frontal lobe; possible right mesial temporal atrophy | TLE | 5 depth 1x16 | RAH5-8 | R Temporal Lobectomy | IB | 22 |
| 3 | 46, M | Focal Impaired Awareness Seizures | Right Hippocampal Sclerosis | TLE | 5 depth 1x16 | RAH4 | R Temporal Lobectomy | IA | 41 |
| 4 | 34, M | Focal Impaired Awareness Seizures | Bilateral hippocampal abnormalities with mild left-sided atrophy | TLE | 2 depth (1x14)<br>3 depth (1x16)<br>2 grid (2x6)<br>1 depth (1x10) | RAM1-2,<br>RHP1-3 | R Temporal Lobectomy | IA | 2 |
| 5 | 57, F | Focal Impaired Awareness Seizures | None | TLE | 11 depth (1x10)<br>1 depth (1x12) | LHH1-3,<br>LTH2-3 | L Lateral Temporal Lobectomy,<br>L Amygdala and | IA | 23 |
|  |  |  |  |  |  |  | Hippocampal<br>Resection |  |  |
| 6 | 53, F | Focal<br>Impaired<br>Awareness<br>Seizures | Left Mesial<br>Temporal<br>Sclerosis | TLE | 10 depth<br>(1x10) | LTH1 | L Temporal<br>Lobectomy | IA | 32 |
| 7 | 54, F | Focal<br>Impaired<br>Awareness<br>Seizures | None | TLE | 10 depth,<br>(1x10) | RHH1-4,<br>RAM1-2 | R Temporal<br>Lobectomy | IB | 26 |

The remaining thirteen patients included in this study were obtained from the freely available online database associated with [14] at iEEG.org (http://crcns.org/data-sets/methods/ieeg-1/about-ieeg-1). These patients underwent invasive EEG recordings with subdural and/or depth electrodes from March 2012 to April 2016 at the University Hospital Zurich as part of their presurgical evaluation. We refer to these thirteen patients as the ETH Zurich dataset. We included all patients that reported good clinical outcomes (class 1) at least 1 year following resective surgery based on the International League Against Epilepsy (ILAE) scale. Information regarding electrode types, data acquisition, and sleep scoring can be found in [14]. Patient demographics for the ETH Zurich dataset are in Table 2.

**Table 2.** Patient demographics of the ETH Zurich dataset. Abbreviations: depth = depth electrode; ETE = extratemporal epilepsy; ILAE = International League Against Epilepsy; Les = lesionectomy; sAHE = selective amygdala hippocampectomy; strip = strip electrode; TLE = temporal lobe epilepsy

| Patient | Age,<br>Gender | Epilepsy | Type of<br>electrodes | Number of<br>channels in<br>resected tissue | Number of<br>channels in non-<br>resected tissue | Surgery | ILAE<br>outcome | Postoperative<br>follow-up<br>(months) |
| --- | --- | --- | --- | --- | --- | --- | --- | --- |
| 8 | 25, M | TLE | 5 depth<br>1 strip 4 x 1<br>1 strip 6 x 1 | 9 | 14 | sAHE;<br>Les | I | 12 |
| 9 | 33, M | TLE | 8 depth | 12 | 12 | sAHE;<br>Les | I | 29 |
| 10 | 20, F | TLE | 5 depth | 12 | 3 | sAHE | I | 13 |
| 11 | 20, F | TLE | 8 depth | 12 | 12 | sAHE | I | 41 |
| 12 | 40, M | TLE | 8 depth | 12 | 12 | sAHE | I | 14 |
| 13 | 48, M | TLE | 8 depth | 12 | 12 | sAHE | I | 11 |
| 14 | 37, M | ETE | 1 grid 8 x 4<br>2 strips 4 x<br>1 | 3 | 31 | Les | I | 36 |
| 15 | 36, M | ETE | 1 grid 8 x 8<br>1 depth | 3 | 62 | Les | I | 37 |
| 16 | 49, M | ETE | 1 grid 8 x 4<br>1 depth | 21 | 16 | Les | I | 25 |
| 17 | 17, M | ETE | 1 grid 8 x 8<br>1 depth | 3 | 50 | Les | I | 25 |
| 18 | 46, F | ETE | 2 grids 8 x 2<br>1 strip 6 x 1<br>1 strip 4 x 1<br>1 depth | 1 | 29 | Les | I | 10 |
| 19 | 31, F | ETE | 1 grid 8 x 4<br>2 strips 4 x<br>1 | 4 | 23 | Les | I | 25 |
| 20 | 17, F | ETE | 1 grid 8 x 4<br>1 depth | 3 | 34 | Les | I | 19 |

### 2.2 iEEG data acquisition

Intracranial EEG was recorded for each patient using a combination of subdural grids and strips, as well as depth electrodes. Recordings from the UCI dataset were collected using a Nihon Kohden JE-120A amplifier with a minimum sampling frequency of 2000 Hz for all patients. Recorded data were re-referenced to a bipolar montage for analysis. All bipolar re-referenced channel pairs that included an SOZ channel were deemed as SOZ; for example, in UCI patient 3, the SOZ was localized to channel RAH4 using the original recording reference, so bipolar channels RAH3-4 and RAH4-5 were considered SOZ for our analysis. Five 3-minute epochs from one night of iEEG recording were randomly selected for each UCI patient, for a total of 15 minutes per patient. Each epoch was chosen from data recorded between 8pm and 8am, as HFO rates are increased during sleep, and the occurrence of muscle artifacts is reduced during slow-wave sleep compared to wakefulness [22–24]. We confirmed that all epochs were interictal data, recorded at least 1 hour away from seizures to reduce the influence of seizures on HFOs [25]. We analyzed all implanted electrodes, which ranged from 80 to 128 contacts for each UCI patient.

All interictal recordings from the ETH Zurich dataset were obtained at least 3 hours away from seizure activity, and we analyzed five 3-minute epochs of slow wave sleep from one night of recording. We included the same channels used for analysis by the original authors; the included and excluded channels can be found in the supplementary information of [14]. Of note, the number of analyzed channels for each patient ranged from 15 to 53. For the analysis in the current study, we will define the “SOZ channels” to be the bipolar re-referenced channels within the resected area for each patient; the specific sites of seizure onset within the resected area were not provided. Defining the SOZ based on the resected volume will likely overestimate the true SOZ in the ETH Zurich dataset, while the method used for the UCI dataset is likely an underestimate due to limited sampling of brain tissue using iEEG electrodes. These datasets are therefore complementary and represent the full range of possible SOZ definitions.

### 2.3 Comparison of SOZ localization using different HFO detection methods

To measure the impact on SOZ localization, we detected HFOs in three different ways. First, we used an automated algorithm with the parameters as originally defined (Sections 2.3.1 and 2.3.2). Second, we visually marked HFOs (Section 2.3.3). Third, we used the same automated algorithm after optimizing the parameters based on the visually marked events (Section 2.3.4).

#### 2.3.1 Automated HFO detection

Automated HFO detection was performed on all iEEG data using an algorithm based on the root-mean-square (RMS) amplitude of bandpass filtered data [26]. We will refer to this as the RMS detector. We used the RMS detector because it is referenced often as a benchmark for new algorithms [17, 20, 27], serves as the core of many other detectors [16, 28-32], and is algorithmically simple to implement and examine. In this algorithm, broadband data is bandpass filtered in the 100-500Hz frequency range, and candidate events are identified when the RMS amplitude of the bandpass filtered signal exceeds a threshold for a minimum duration (which we will abbreviate as “min_dur;” see Fig. 1a). The RMS signal is calculated using a moving window (with the duration termed “RMS_win”), and the first threshold (“nSD1”) is defined as five standard deviations above the mean RMS signal. Consecutive events separated by less than a predefined gap time (“gap”) are combined into one event. Candidate events are retained when a minimum number of peaks (“min_pk”) in the rectified filtered data exceed a second threshold (“nSD2”) defined as 3 standard deviations above the mean rectified data.

**Fig. 1.**
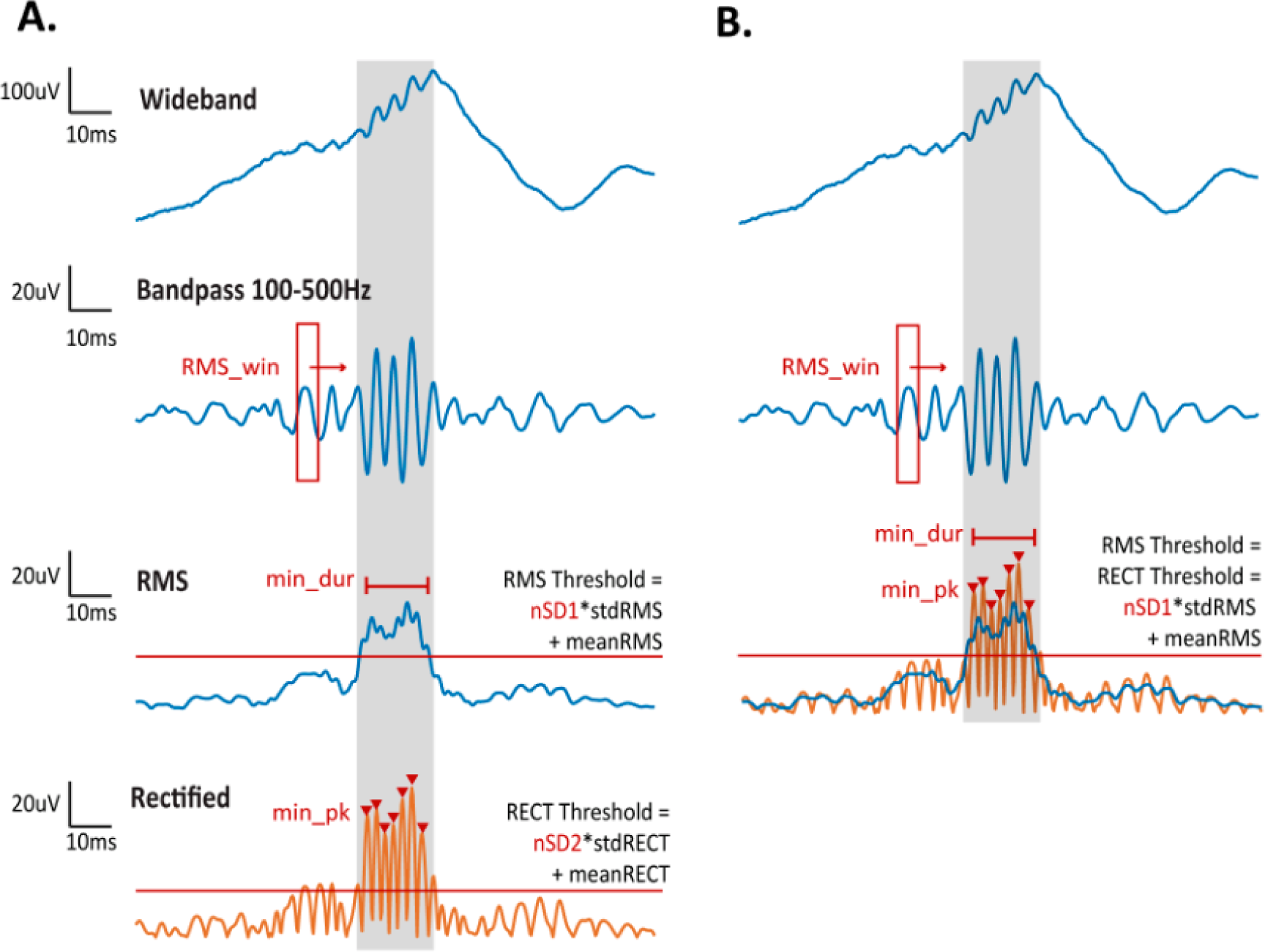
Automatic HFO detection algorithm where **A.** is the Full RMS detector and **B.** is the Reduced RMS detector used in our analysis. The shaded gray region represents the window containing the identified candidate event. Detection thresholds are indicated by horizontal red lines, and each peak in the candidate event is marked by a red triangle. Abbreviations: RMS = root-mean-square, std = standard deviation

To reduce the algorithm complexity, we set threshold two equal to threshold one (nSD2 = nSD1), as they both ensure that the signal’s energy exceeds a threshold determined from baseline activity (Fig. 1b). As in the original algorithm, we required that a minimum number of 6 peaks in the rectified filtered data exceed this threshold, to promote rejection of fast transients. All other steps in the original algorithm were maintained. The detection algorithm used in our analysis thus contains five parameters: RMS window size (RMS_win), minimum event duration (min_dur), number of standard deviations above the mean RMS signal (nSD1), minimum gap time (gap), and minimum number of peaks (min_pk). From these parameters, we varied the three that directly affect initial detection of candidate events: RMS window size, minimum event duration, and threshold. Because the minimum gap time is a post-processing step to join candidate events and the minimum number of peaks has little impact as a criterion due to the redundancy with minimum event duration, we kept these variables constant at their default values. The impact of minimum duration on SOZ localization was tested by varying its value from 6 ms to 12 ms. We found no significant difference in localization accuracy for the two values, and we therefore used a fixed value of 6 ms for this study (Supplementary Figure 1). The default value of each parameter and the ranges of values tested are listed in Table 3.

**Table 3.** Detection parameters varied and corresponding ranges of values. Default values are underlined. The default value for minimum gap time is 10 ms and minimum number of oscillations is 6.

| Parameter | Values |
| --- | --- |
| Threshold (nSD1) | 1, 2, 3, 4, <u>5</u>, 6, 7, 8, 9, 10, 11, 12, 13, 14, 15 |
| RMS window size (RMS_win) (ms) | 2, <u>3</u>, 5, 7, 9, 11, 15, 17, 20 |
| Minimum event duration (min_dur) (ms) | <u>6</u>, 12 |

#### 2.3.2 Automatic Artifact Rejection

Because the RMS detector is highly sensitive [17], we implemented two artifact rejection methods to improve specificity. Both methods, PopDet and BkgStabaDet, were introduced by [31] for the purpose of creating a generalized HFO detection algorithm for long-term intracranial EEG recordings, such that the algorithm automatically identifies quality HFOs without any patient-specific tuning or operator intervention [31]. These artifact rejection steps were designed using the RMS detector for initial detection, making them appropriate for our analysis. In this study, the default parameters from [31] were used for all patients and all parameter sets.

The PopDet criterion was designed to detect DC shifts and fast transients commonly found in EEG data. When a DC shift or fast transient is filtered, it can have the appearance of an HFO in the 100-500 Hz frequency band [32, 33]. However, these transients also contain power at very high frequencies, whereas a true HFO should have band-limited power. Therefore, PopDet identifies instances when the line length of a 0.1s window in the 850-990 Hz frequency range exceeds a threshold of 5 standard deviations above the mean line length calculated from baseline. Baseline is defined as a 5-second window preceding the window being evaluated.

The BkgStabaDet was designed to detect spatially diffuse HFOs, which are considered false positive detections because they contradict the idea that HFOs should be focal events [34, 35]. If an HFO occurs in all channels of a depth electrode or grid, it will appear in the common average. Therefore, detections in a single channel that occurred within 100ms of a detection in the common average were excluded from further analysis. As in [31], we applied the RMS detector from [26] to the common average within each depth electrode or grid using the default parameters.

To assess the impact of the automated artifact rejection steps, we calculated the percentage of candidate HFOs rejected for each parameter set and the SOZ localization accuracy before and after artifact removal. The percentage of rejected candidates by PopDet was calculated as the total number of candidates marked by PopDet over the total number of candidates counted across all epochs. The same procedure was applied to candidates marked by BkgStabaDet. Because a candidate can be marked by both PopDet and BkgStabaDet, the total percentage of rejected candidates by PopDet and BkgStabaDet can exceed 100. Overall, we found that the percentage of rejected candidates was low, except for detectors using the highest values of nSD1 and the lowest values of RMS_win (a representative example is shown in Supplementary Figure 2). Moreover, we found that artifact removal did not significantly impact SOZ localization accuracy (Supplementary Figure 2). However, these processing steps were retained for our analysis, as rejection of false positive detections is standard practice.

#### 2.3.3 Visual Marking of HFOs

Visual marking of HFOs was completed by three reviewers, with each reviewer marking a nonoverlapping subset of the 20 patients. Each reviewer manually marked HFOs in one minute of iEEG data from each patient. A custom graphical user interface displayed 600 ms of the broadband data for eight iEEG channels, as well as the corresponding data filtered in the ripple band (100-500Hz), and the reviewer could move forward/backward in time in 60 ms intervals. The broadband data were used to confirm there were no artifact events, sharp transients, or DC shifts that may appear as false HFOs in the filtered data.

The reviewer could select whether they had high confidence or low confidence that each event was an HFO. Events were considered high confidence if the amplitude of the event stood out from the surrounding background for four or more oscillations; this event could not be interrupted by more than one low-amplitude oscillation. Low confidence events typically had one feature that did not match the prototypical HFO. For example, there were cases of borderline high amplitude (especially when the amplitude of the background changed over time and was difficult to estimate), and sometimes the high amplitude oscillations were interrupted for a short period of time or had an irregular amplitude envelope. Events occurring simultaneously in a large number of channels were considered artifactual because we expect HFOs to be local events. For this study, we analyzed only high confidence detections, as those events fit the full, typical definition of an HFO.

#### 2.3.4 Optimizing HFO detection settings to match visual marking

To determine the HFO detection settings that best matched the visually marked events, we calculated the number of automatically detected events that overlapped with the manually marked events in the same segment of iEEG. More specifically, visual and automatic detections were considered overlapping if 50% of the shorter detection overlapped with the longer detection. Events that were marked both manually and by the detector were considered true positives (TP), events marked manually but not by the detector were classified as false negatives (FN), and events marked only by the detector were false positives (FP). Due to the lack of a true negative, the F1 score was used to identify the parameter set that most closely matched the manual markings:

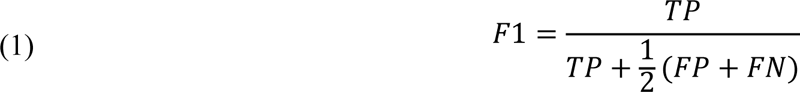

The F1 score was calculated for all combinations of RMS_win and nSD1 values used for automatic detection. The parameter set with the highest F1 score was the “optimal” set based on visual markings. If there were multiple parameter sets with the same F1 score, we selected the parameter set with the lowest nSD1 and the highest RMS_win, as this set should have the highest sensitivity.

### 2.4 Measurement of localization accuracy

We used a receiver operating characteristic (ROC) curve to quantify the ability of HFO rate to localize the clinically determined SOZ. For channels labeled SOZ, those with HFO rates that exceeded the threshold were marked as true positives (TP), and those that did not exceed the rate threshold were marked as false negatives (FN). For channels defined clinically as non-SOZ (nSOZ), those that had HFO rates below the threshold were marked as true negatives (TN), and those with HFO rates above the threshold were marked as false positives (FP). The ROC curve was plotted by varying the HFO rate threshold and determining the true positive rate (TPR) and false positive rate (FPR) for each value:

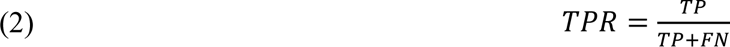

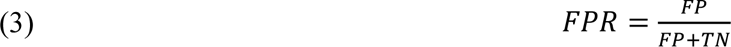

The area under the ROC curve (which we will abbreviate AUC) was determined for all patients and HFO detection methods.

As a complementary measure, Precision-Recall (PR) scores were evaluated to examine the positive predictive value for each parameter set. Precision-Recall analyses are preferred when prediction power of imbalanced classes is being evaluated; in our case, we have imbalanced classes represented by the small subset of SOZ channels compared to the larger class of nSOZ channels for UCI patients, and the small subset of nSOZ channels compared to the larger class of SOZ channels for some ETH Zurich patients (specifically, patients 10 and 16). Precision (P), Recall (R), and the F1 score (F1) were computed as follows:

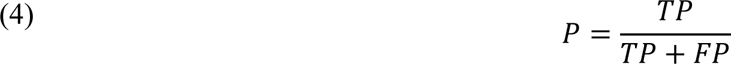

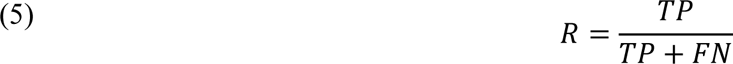

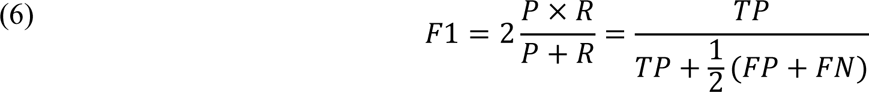

Like the ROC curve, the PR curve was constructed by varying the rate threshold and calculating the precision, recall, and F1 score for each value. Then the AUC of the ROC curve and maximal F1 score of the PR curve were determined for each detection method (visual, automatic, and optimized automatic) and for each pair of parameters for the automatic approaches.

## 3. Results

### 3.1 Parameter Selection using visually marked events does not improve SOZ localization

Consistent with prior studies, our data showed an association between high HFO rates and the SOZ (Figure 2). 15 out of the 20 patients had a maximal F1 score greater than 0.5 for at least one of the detection methods. Similarly, 18 patients had AUC scores greater than 0.7, suggesting that HFOs were associated with the SOZ in most patients. Also consistent with prior studies, there were a few patients with low maximal F1 and AUC scores, indicating that sometimes the recorded HFOs were not sufficient for identifying the SOZ.

**Figure 2.**
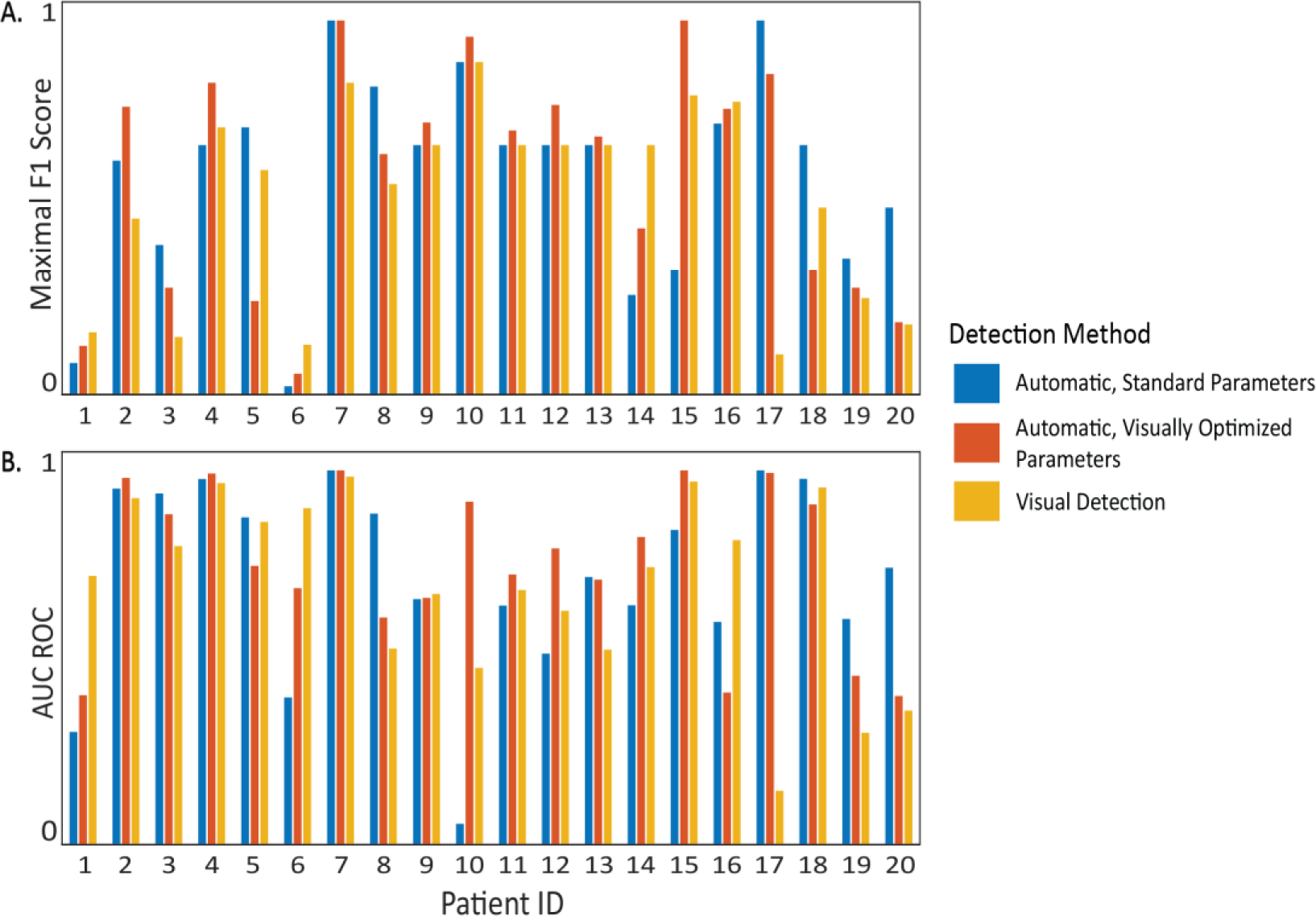
SOZ localization accuracy based on **A.** the maximal F1 score, and **B.** the AUC of the ROC for the three HFO detection methods.

While overall SOZ localization accuracy varied across patients, we found that the results using the three detection methods were comparable. When comparing pairs of HFO detection methods, there were no statistically significant differences for either the AUC or the maximal F1 scores across patients (Table 4). Based on both the AUC and the maximal F1 score, the highest accuracy was obtained for visual marking in three patients (1,6,16), the visually optimized parameter settings in six patients (2,4,10-12,15) and the standard parameter settings in seven patients (3, 5, 8, 17, 18, 19, 20). Across patients, there was a correlation between the maximal F1 scores for all pairs of methods, with the highest correlation occurring between visual detection and automated detection with visually optimized parameters (Table 4).

**Table 4.** The three methods of HFO detection produced comparable SOZ localization results. Using the Wilcoxon signed rank test, p-values indicate that there are no significant differences when comparing the AUC values (column 2) and maximal F1 score (column 3) between methods. The last column contains the Pearson correlation values between detection methods, along with their corresponding p-values.

| Detection Methods | AUC of ROC | Maximal F1 | Correlation of Maximal F1 (p value) |
| --- | --- | --- | --- |
| Automatic Standard vs Automatic Visually Optimized | p=0.7172 | p=1 | 0.677 (p<0.05) |
| Automatic Standard vs Visual | p=0.9702 | p=0.2238 | 0.461 (p<0.05) |
| **Automatic Visually Optimized vs Visual** | $p=0.2043$ | $p=0.1506$ | $0.712 (p<0.05)$ |

### 3.2 Visually Optimized Parameters vary across patients

The visually optimized detection parameters were not consistent across patients and did not appear to be related to the overall SOZ localization accuracy. For most patients, the thresholds were lower and the RMS windows were longer than the standard parameters (Figure 3). There were three outlier patients for which the RMS window had its maximum duration (20 ms), but there were no obvious commonalities between the subjects. The localization accuracies for these subjects were not notably high nor low. Furthermore, these patients had HFOs marked by different reviewers and originated from both the UCI and ETH Zurich datasets. One important factor to consider when interpreting these results is that the overlap between visually detected events and automatically detected events was generally low. F1 scores for comparing the two types of events (and ultimately selecting the automatic detection parameters) had a median value of 0.347 and interquartile range of 0.186, suggesting high amounts of false positives and/or false negatives.

**Figure 3.**
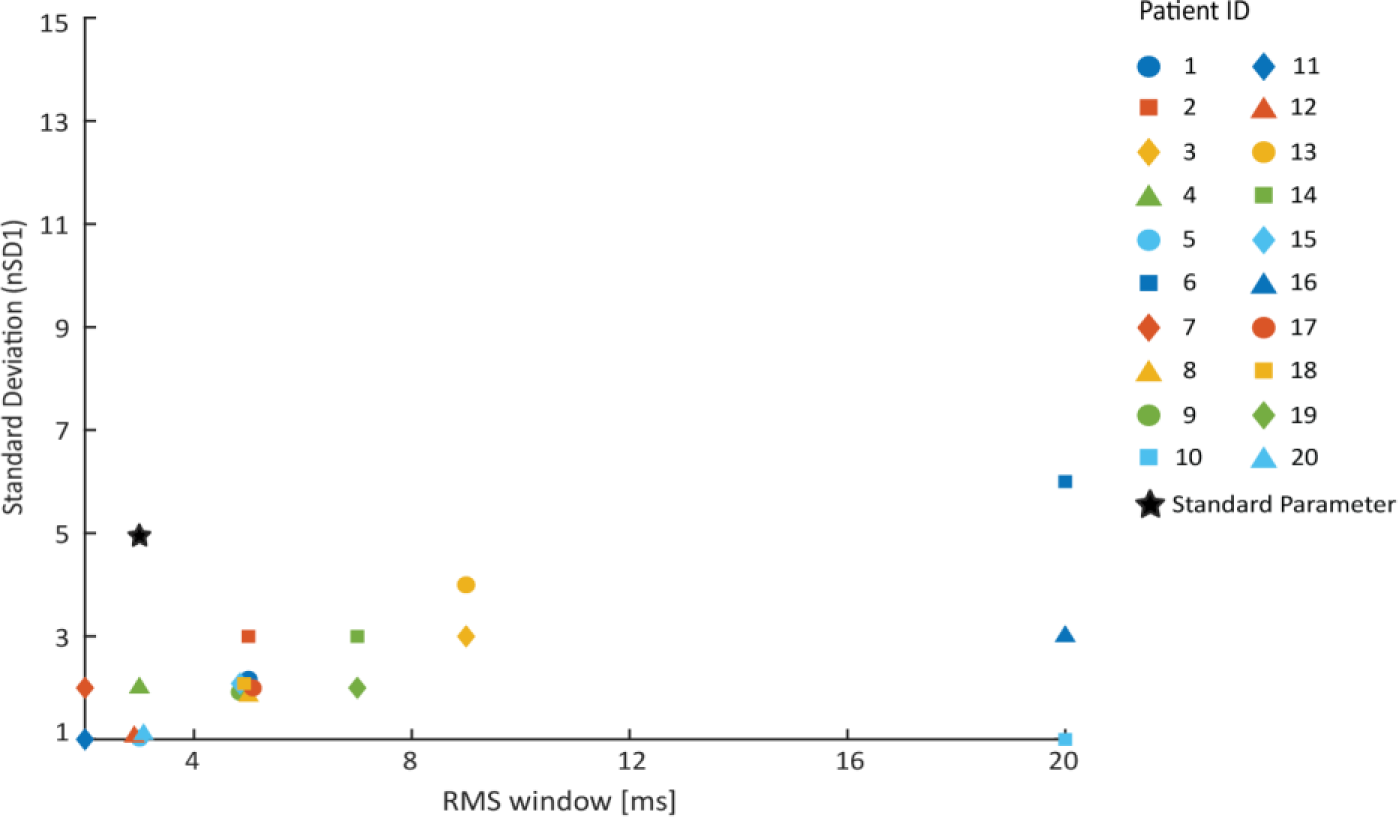
Distribution of visually optimized detector settings. Compared to the standard detection settings (black star), many of the optimized parameter settings had lower thresholds (nSD1) and longer RMS window lengths.

### 3.3 Regions of high localization accuracy are patient-specific and vary across a wide range of parameters

To provide context for the SOZ localization accuracy using the three different HFO detection strategies, we measured the maximal F1 score and AUC of the ROC for a range of automatic detection parameters in each subject (Figure 4). Specifically, we varied RMS_win from 2 to 20 ms, and nSD1 ranged from 1 to 15. A band of generally high accuracy was found for most patients as the RMS window and threshold increased together (indicated by the red bands stretching from the lower left corner to the upper right corner). In many subjects, this band included all RMS window lengths, indicating that this parameter had a low overall influence on localization accuracy. This band of high localization accuracy was frequently seen in plots of the AUC and often spanned more than half of the tested thresholds as the RMS window length increased, e.g., patients 2-5, 7, 8, 13, 17, 18 in Figure 4. The maximal F1 was more stringent, with high localization accuracies often occurring for a smaller range of thresholds, e.g., patients 4, 5, 15. The same diagonal pattern could even be seen in patients with poor localization performance, e.g., patients 1 and 6.

**Figure 4.**
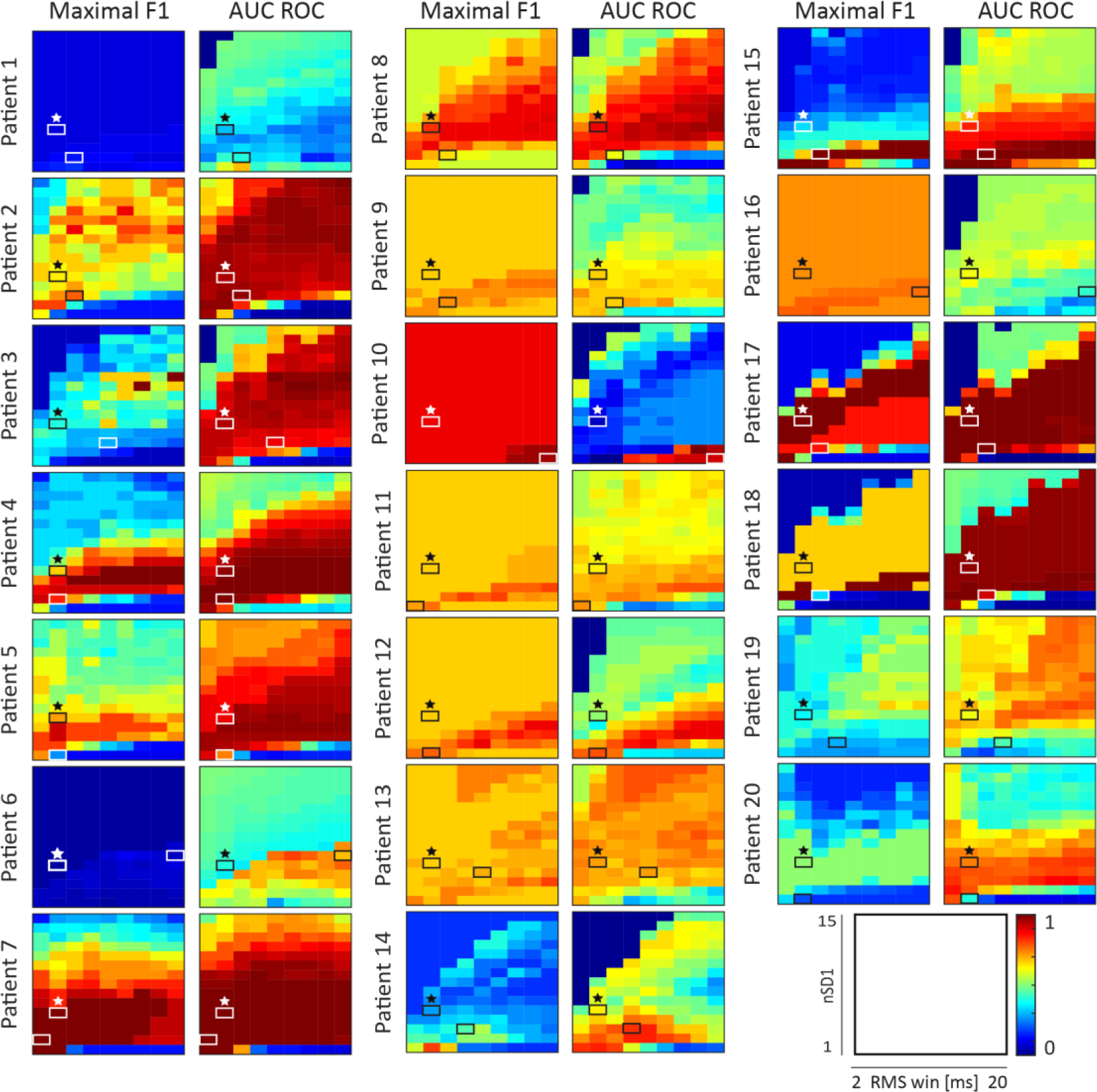
Results of SOZ localization accuracy for all automated HFO detection settings. For each patient, the maximal F1 scores (left) and AUC values (right) are shown for all pairs of parameter settings. Starred boxes (black or white, chosen to maximize contrast) indicate the SOZ localization accuracy for the standard detection settings. The second box (black or white) in each subfigure indicates the visually optimized detection settings for each patient.

As we know from Figure 2, some patients had high SOZ localization accuracy using standard HFO detection settings, while others did not. Several patients showed localized regions of high accuracy that differed from the standard parameter settings, e.g., patients 3, 10, 12, 14, 15, and 19 in Figure 4. Optimization based on visually marked HFOs assisted in identifying these regions of high accuracy in some patients (e.g., patients 9, 10, 12, 14, 15), while making localization accuracy worse in others (e.g., patients 3, 8, 19 and 20).

### 3.4 Class imbalance affects interpretation of PR and ROC Results

Comparison of the maximal F1 score and the AUC within individual subjects can also provide important context for the results. For all seven patients from the UCI dataset (patients 1-7) and 6 patients from the ETH Zurich dataset (14, 15, 17-20), the ROC provided a greater range of detection parameters with higher localization accuracies than the PR results [36]. This can be explained by the larger proportion of non-SOZ to SOZ channels, which affects the precision and false positive rate measures used to generate the PR and ROC curves. Precision is determined by the number of true positives relative to all predicted positives, as defined in equation 3. In patients where the number of nSOZ channels considerably outnumbers the SOZ channels, there is an increased probability of marking a false positive, which increases the denominator of equation 3 and reduces the precision. Furthermore, because there are significantly fewer SOZ channels, there are fewer true positives for both the precision and recall equations, leading to lower maximal F1 scores. In contrast, the false positive rate in ROC analysis is interpreted as the proportion of nSOZ channels with HFO rates above the threshold. In patients with a large number of nSOZ channels (and thereby a high likelihood of having true negative channels based on HFO rate), the false positive rate can generally be kept low while the true positive rate increases, resulting in more favorable ROC results. An example of this is shown for patient 6 in Supplementary Figure 3.

In contrast, seven patients from the ETH Zurich dataset (patients 8-13, 16) had larger regions of high localization accuracy in the PR results (Fig. 4, left column) compared to the ROC results (Fig. 4, right column). This relationship is inverted compared to the other subjects due to the smaller number of electrodes and broader interpretation of the SOZ, which was defined by the resected volume. These patients had a similar number of SOZ and nSOZ channels, improving the likelihood of obtaining favorable positive predictive performance. Furthermore, due to the decreased number of electrodes, we see less variation in F1 score across the parameter space for most of these ETH Zurich patients.

## 4. Discussion

Our study set out to understand the impact of HFO detector settings and optimization on SOZ localization accuracy. SOZ localization performance was comparable across the tested detection methods, suggesting that using standard parameter settings is acceptable for HFO detection. On the other hand, when exploring the HFO detection parameter space to provide context for this result, we found that the regions of high SOZ localization accuracy were patient-specific. This result suggests that, while the standard HFO detection settings may perform adequately at the group level, the use of individualized settings has the potential to improve SOZ localization accuracy. However, optimization using visually marked events did not enable selection of the parameter settings with the highest localization accuracy in each patient. Therefore, there is a need for novel approaches to this challenging problem.

One difficulty we encountered in our analysis was the inability to find metrics that could robustly compare visually marked events to automatically detected HFOs and quantify SOZ localization. First, the F1 score was used to quantify the overlap between visual and automated detection of events, as it does not require true negatives. Previous studies used selected baseline periods without any oscillatory activity to calculate the true negative rate [20], however this method would have further added to the high burden of visually marking HFOs in our study. Furthermore, such baseline periods can be marked in a highly selective way, such that they would never be falsely detected, and if the true negative rate is always perfect, its utility is diluted. However, the F1 score was not ideal, as evidenced by its generally low values. Automatic detection using low thresholds and short RMS windows generally resulted in an HFO rate comparable to visual detection and therefore a balanced number of true positives, false positives, and false negatives. For this reason, our optimized parameters tended towards the lower left corner of the parameter space (Figure 3). However, the visual markings were highly specific, which resulted in low true positive values and low F1 scores. As an alternative to the maximal F1 score, we tested the positive predictive value (PPV), but this metric also biased the results toward certain regions in the parameter space. Because PPV relies only on true positives and false positives, the optimal parameters had higher thresholds, where there were very few automatic detections and thus low false positive rates. Thus, the use of PPV instead of the maximal F1 score did not improve the seizure localization accuracy for this cohort of patients (data not shown). Second, when measuring SOZ localization accuracy, we found very different results for the maximal F1 score and the AUC (Figure 4). The AUC of the ROC often showed a wider range of parameters with higher accuracies, but this required careful interpretation based on the balance of SOZ and nSOZ channels in each patient. Therefore, we used both measures, as they are complementary and provided a more holistic view of the results.

The data selection and visual marking of HFOs are potential sources of variability in this study. For visual detection of HFOs, we implemented an iEEG review method that would be practical in a clinical setting, in which a single, trained reviewer marked data from each patient. Manual markings of events can be confounded by persistent high amplitude background activity, frequent artifacts (which could cause the reviewer to overlook or dismiss events in that channel), or physiological HFOs generated from healthy tissues [37]. An example of this was seen in patient 17, where the SOZ contained extremely high amplitude events that shared characteristics with HFOs, making it difficult to distinguish true HFO events. Furthermore, it is possible that the randomly selected iEEG segments were not representative of the patients’ long-term brain activity, as HFO rates are known to fluctuate over time [38, 39]. This could have impacted both the visual markings and automated detection. For example, patients 3 and 10 in our study showed low rates for manual and automatic methods, respectively. It is possible that this was specific to the selected iEEG segments. Random selection of segments is a rigorous approach and is easily replicated in a clinical environment, but the use of a larger number of segments would ensure that the results are representative for each patient.

Concerns about bias in visual marking were mitigated by the fact that the SOZ localization accuracies across all three detection methods were highly correlated. In particular, the results for visual marking were significantly correlated to automatic detection with standard parameters (Table 4). This suggests that the visually marked events were relatively consistent with those resulting from an objective, independent algorithm and not overly impacted by individual bias.

## Conclusion

Our results demonstrate that SOZ localization using HFOs is relatively robust to the method of HFO detection when the same method is used for all patients. This suggests that automated HFO detection using standard parameters is an acceptable approach; this is advantageous, as it eliminates the need for time consuming manual marking of events. However, our results also suggested that individualized optimization of HFO detection has the potential to significantly improve identification of the SOZ for individual patients. If novel methods can be developed to accurately identify these optimal parameters, this would improve the accuracy of HFOs as a biomarker of the SOZ. Ultimately, this would allow clinicians to identify and resect epileptogenic tissue with greater precision, thus leading to higher rates of seizure freedom after surgery.

## Supplementary

**Supplementary Figure 1.**
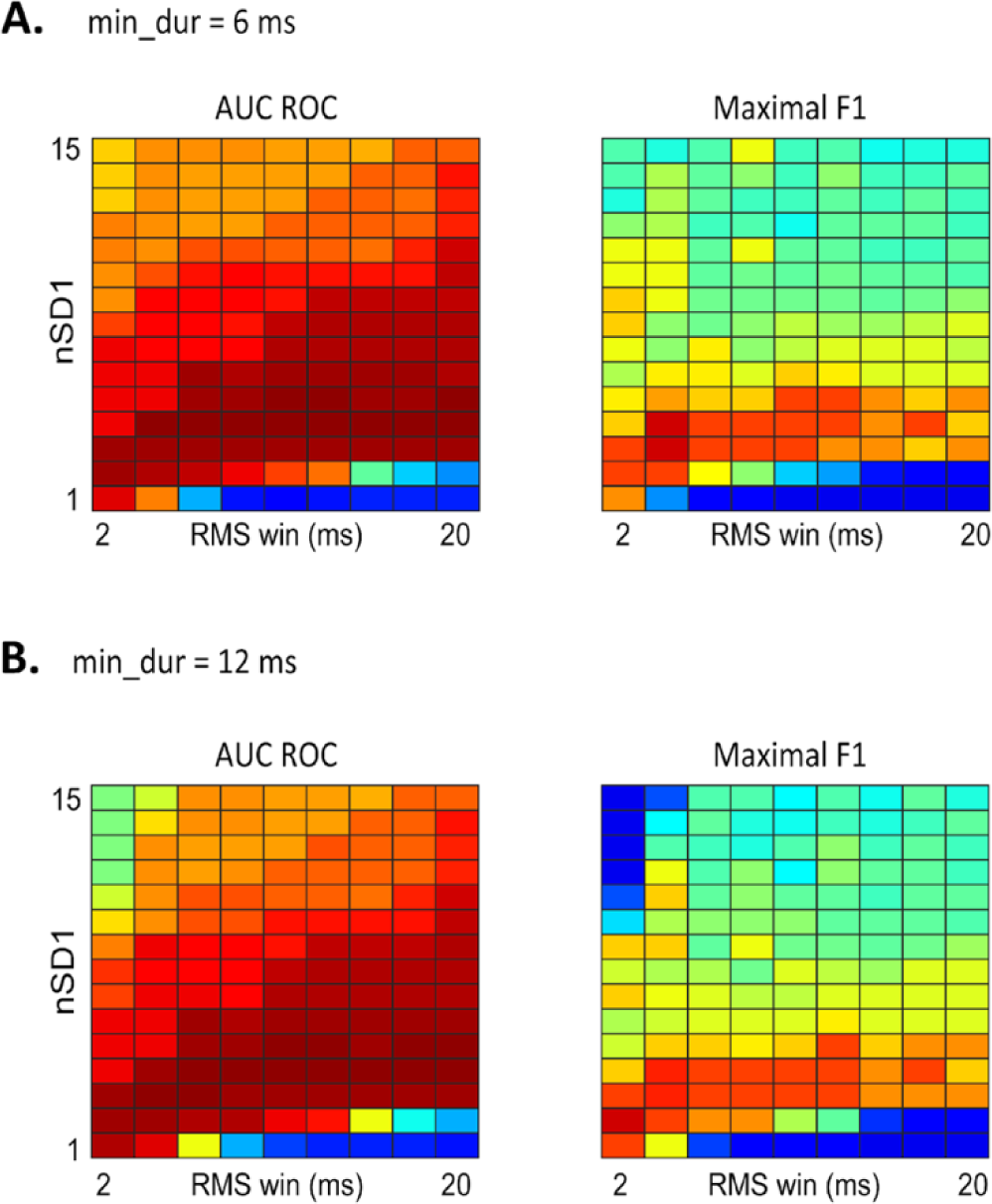
Minimum event duration does not affect SOZ localization performance. Representative examples of SOZ localization performance across the parameter space, comparing results calculated using minimum event duration of **a** min_dur=6ms and **b** min_dur=12ms. Heatmaps of AUC of the ROC curve and maximal F1 score are shown across the parameter space for patient 5, with RMS window length (RMS_win) varying on the horizontal axis and RMS threshold (nSD1) varying on the vertical axis. All values range from 0 (dark blue) to 1 (dark red), with results closer to 1 indicating good classification of SOZ and nSOZ channels

**Supplementary Figure 2.**
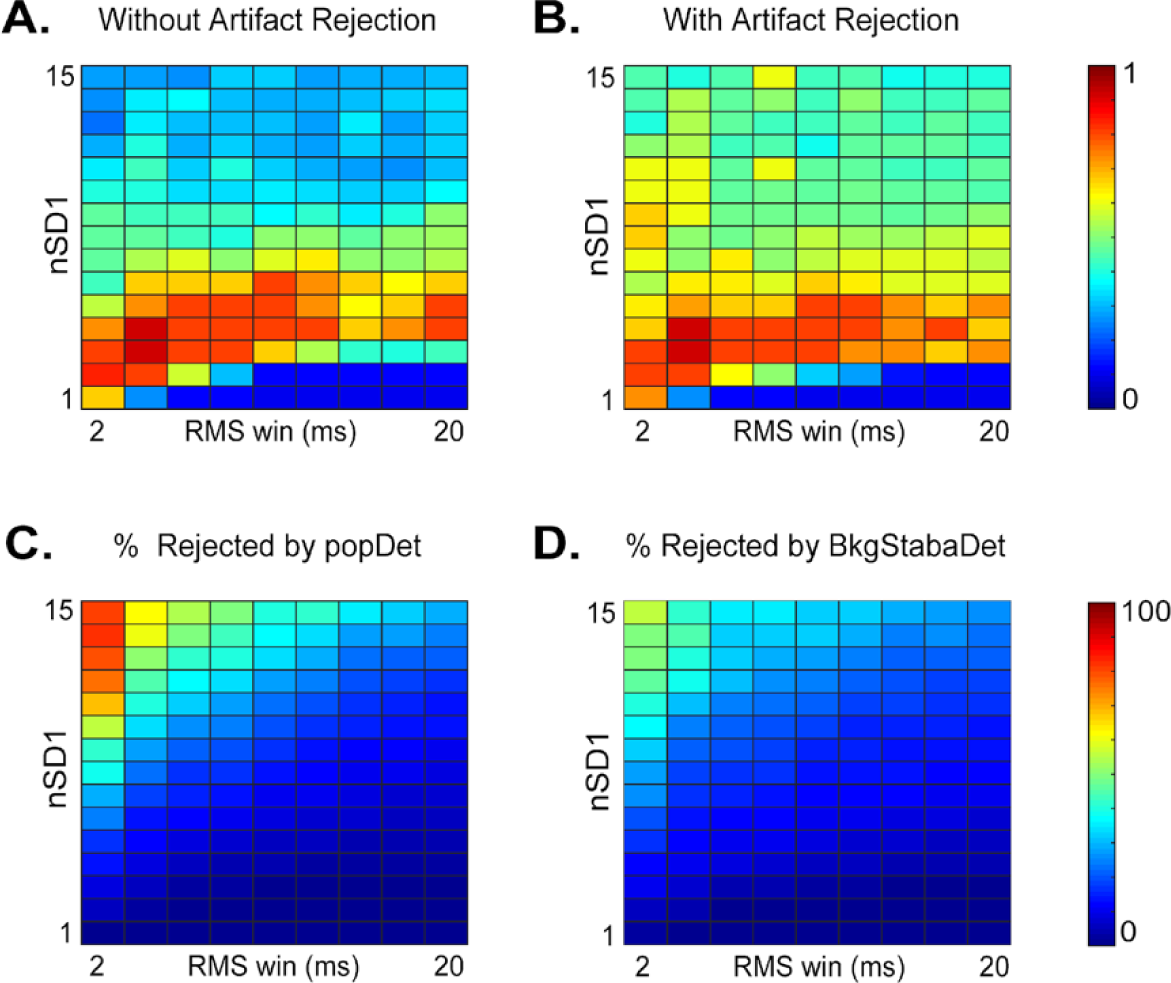
Artifact rejection does not impact parameters with higher SOZ localization accuracy. Comparison of maximal F1 scores **a** without and **b** with artifact rejection across the parameter space. Percentage of candidate events rejected by **c** PopDet and **d** BkgStabaDet across the parameter space. Results from patient 5 are shown as a representative example.

**Supplementary Figure. 3.**
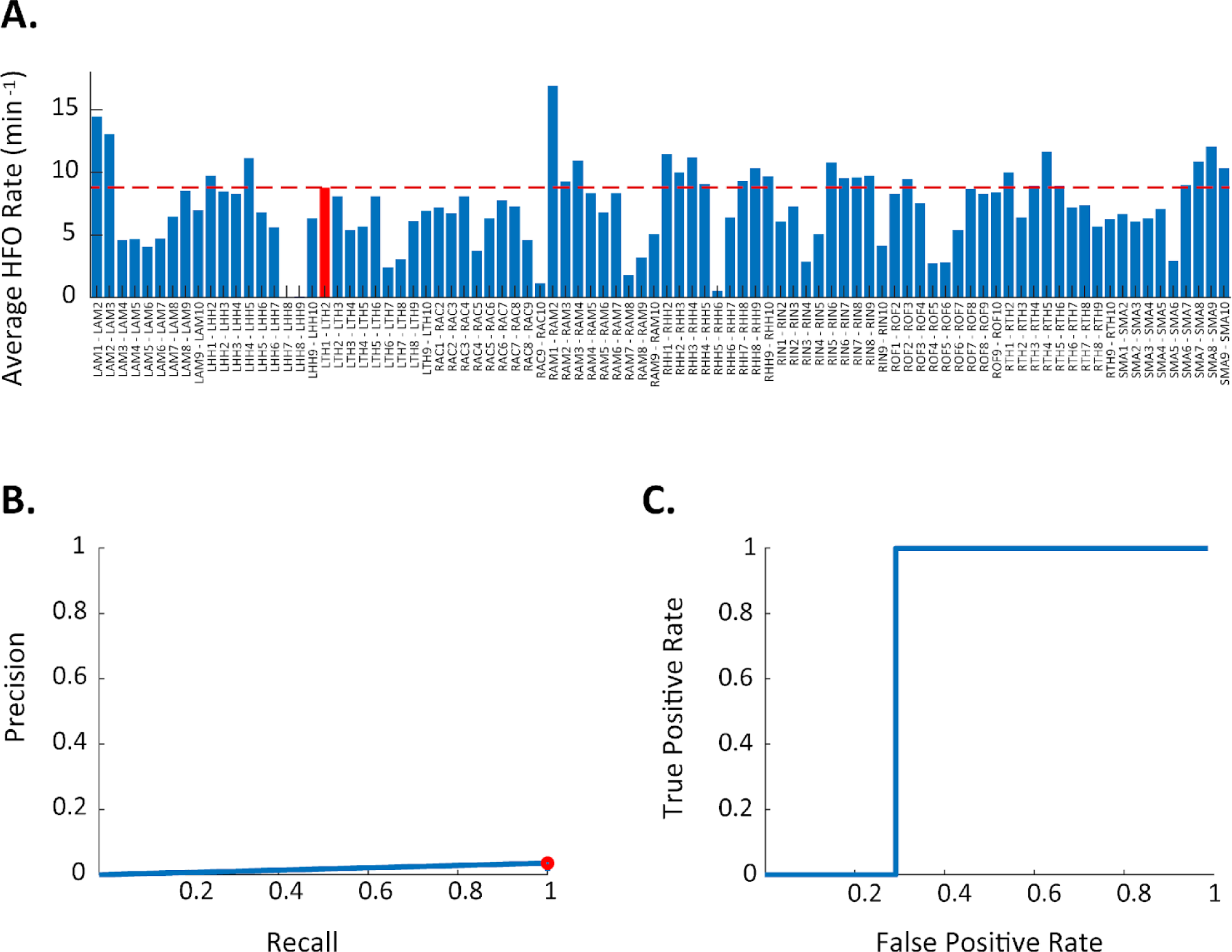
Imbalanced proportion of SOZ channels and nSOZ channels alters interpretations of precision-recall and receiver-operator-characteristic results. **a** Average HFO rate for each channel in Patient 6 using an RMS threshold of three standard deviations (nSD1=3), RMS window size of 20ms (rms_win=20ms), and minimum event duration of 6ms (min_dur=6ms). In this case, we observe that the HFO rate distribution contains a considerable number of nSOZ channels with high HFO rates. **b** The resulting PR curve with the maximal F1 point circled in red, where Twenty-four nSOZ channels have higher HFO rates than the SOZ channel, thereby resulting in a poor PR curve. **c** The ROC curve shows better results due to the large proportion of nSOZ channels with lower rates compared to the SOZ channel. The low number of SOZ channels (relative to the large number of nSOZ channels) negatively impacts the PR curve when the number of false positives is high, but it minimally impacts the ROC curve.

